# Examining the occupancy-density relationship for a low density carnivore

**DOI:** 10.1101/066662

**Authors:** Daniel W. Linden, Angela K. Fuller, J. Andrew Royle, Matthew P. Hare

## Abstract

1. The challenges associated with monitoring low-density carnivores across large landscapes have limited the ability to implement and evaluate conservation and management strategies for such species. Noninvasive sampling techniques and advanced statistical approaches have alleviated some of these challenges and can even allow for spatially explicit estimates of density, arguably the most valuable wildlife monitoring tool.
2. For some species, individual identification comes at no cost when unique attributes (e.g., pelage patterns) can be discerned with remote cameras, while other species require viable genetic material and expensive lab processing for individual assignment. Prohibitive costs may still force monitoring efforts to use species distribution or occupancy as a surrogate for density, which may not be appropriate under many conditions.
3. Here, we used a large-scale monitoring study of fisher *Pekania pennanti* to evaluate the effectiveness of occupancy as an approximation to density, particularly for informing harvest management decisions. We used a combination of remote cameras and baited hair snares during 2013–2015 to sample across a 70,096 km^2^ region of western New York, USA. We fit occupancy and Royle-Nichols models to species detection-nondetection data collected by cameras, and spatial capture-recapture models to individual encounter data obtained by genotyped hair samples.
4. We found a close relationship between grid-cell estimates of fisher state variables from the models using detection-nondetection data and those from the SCR model, likely due to informative spatial covariates across a large landscape extent and a grid cell resolution that worked well with the movement ecology of the species. Spatially-explicit management recommendations for fisher were similar across models. We discuss design-based approaches to occupancy studies that can improve approximations to density.

## Introduction

Species distribution and abundance are fundamental quantities in ecology, and serve as the primary state variables for informing large-scale conservation and management (Jones 2011). The choice of which state variable to use as a monitoring tool depends on the types of population inferences regarding variation over time or space needed to meet objectives (Yoccoz, Nichols & Boulinier 2001). In practice, the choice is also dictated by logistical constraints. Noninvasive survey methods have greatly expanded our capacity to monitor certain wildlife species at large scales (e.g., terrestrial carnivores; Long *et al*. 2008), yet the observations required to estimate abundance as opposed to occurrence can still be more costly and difficult to obtain (Burton *et al*. 2015). Thus, monitoring programs may use occurrence as a surrogate for abundance or density (MacKenzie *et al*. 2006; Ellis, Ivan & Schwartz 2014), under the assumption that monitoring objectives can still be achieved.

Relationships between occurrence and abundance have been demonstrated within and between species at macro-ecological scales, with biological and statistical mechanisms used to explain variation in the strength of such relationships (Brown 1984; Gaston *et al*. 2000). From a statistical standpoint, a relationship between species occurrence and abundance should result from the fact that both quantities represent areal summaries of the same spatial point pattern of individuals on a landscape (Kéry & Royle 2016, pg. 3). The summaries are equivalent when the grain (i.e., size of the spatial unit of observation; Wiens 1989) over which the point pattern gets summarized is small enough such that the maximum number of individuals within a unit is 1. Conversely, a large grain which results in most units being occupied with ≥1 individual will produce an occurrence pattern that exhibits no useful relationship with variation in abundance. Regardless of grain, the intensity and spatial variation in the point pattern will also determine how well any summaries of occurrence and abundance match. In wildlife studies, the true point pattern of individuals is almost never known and must be sampled, accounting for both spatial variation and detectability (Pollock *et al*. 2002). Thus, the choice of grain will be constrained by study objectives, species ecology, and possible sampling and analytical frameworks (Wiens 1989). In continuous landscapes without naturally defined spatial units, this decision can be especially complicated and have consequences for the inferences derived from sampling. For example, the sampling unit definition in an occupancy model (sensu MacKenzie *et al*. 2002) will dictate the necessary data collection, affect model assumptions and interpretation, and potentially alter the relationship between estimated occurrence and true population density (Efford & Dawson 2012).

One strategy for selecting the grain in an occupancy study for highly mobile species has been to use previous estimates of home range size as a minimum bound to avoid violating the “closure” assumption (Karanth *et al*. 2011; O’Connell & Bailey 2011). Closure in this case relates to changes in the occupancy state between surveys at a given site due to animal movement, which alters the interpretation of occupancy to mean “use” (MacKenzie & Royle 2005) and differentiates instantaneous from asymptotic occupancy (Efford & Dawson 2012). Selecting a relatively large grain to accommodate wide-ranging, mobile species where movement between surveys is most problematic makes it nearly impossible to truly survey the entire sampling unit (Efford & Dawson 2012). Common noninvasive survey techniques, such as remote cameras, can have very small sampling “footprints” compared to the movements of target species (Clare, Anderson & Macfarland 2015). Even for baited detectors which result in higher observation rates due to larger effective trapping areas at each site (du Preez, Loveridge & Macdonald 2014), the actual trapping area for a given site is still mostly unknown. Finally, selecting the grain to accommodate species ecology becomes complex when movement behavior is sex specific (Sollmann *et al*. 2011), causing some model assumptions to be violated by individuals of one or the other sex. Without understanding how these sampling tradeoffs result in modified observation processes, the interpretation of what occupancy estimates represent may be far removed from the truth, reducing the value of occupancy modeling as a proxy for abundance (Efford & Dawson 2012).

Validation and calibration are important steps in determining the utility of a proxy as a tool for natural resource management (Stephens *et al*. 2015). Previous studies have demonstrated the statistical relationships between species occupancy and density by simulating individual point patterns and hypothetical detection surveys (Efford & Dawson 2012; Ellis, Ivan & Schwartz 2014), providing guidance for the sampling design of large-scale monitoring studies. These types of simulations and power analyses require previous information on species ecology that may not always exist, particularly for widespread species with regional variation. Clare, Anderson and Macfarland (2015) empirically estimated the point pattern of individuals using spatial capture-recapture (SCR) modeling (Borchers & Efford 2008; Royle & Young 2008), and demonstrated a strong relationship between estimates of bobcat *Lynx rufus* occupancy and density using species detections and individual encounters, respectively, collected from remote camera traps. Given the typical capture-recapture requirement of encounter data from identified individuals (but see Chandler & Royle 2013), non-invasive sampling applications of SCR have been limited to species with unique features that can be photographed or to surveys that can collect genetic samples for genotyping (Royle *et al*. 2014). For species without identifiable features or monitoring programs that cannot afford long-term investment in expensive genetic sample processing, a calibration of the occupancy-density relationship could serve to guide monitoring design.

Here, we used a large-scale monitoring study of fisher *Pekania pennanti* to evaluate the effectiveness of occupancy as an approximation to density, particularly for informing harvest management decisions. A medium-sized carnivore traditionally valued for its fur, fisher had been extirpated from much of eastern North America by the early 20^th^ century due to unregulated trapping and habitat loss; recent population expansions have coincided with furbearer protection measures and conversion of farmland to forest in the region (Lancaster, Bowman & Pond 2008). An increased interest in expanding harvest opportunities prompted the New York State Department of Environmental Conservation (NYSDEC) to implement a monitoring program for fisher to identify the wildlife management units that could sustain regulated trapping (Fuller, Linden & Royle 2016). We sampled a large landscape across western New York, USA using baited camera and hair snare traps and fit occupancy models to species detection-nondetection data and spatial capture-recapture models to individual encounter data obtained by genotyped hair samples. We also used the species detection-nondetection data to estimate abundance (density) with the Royle-Nichols model (Royle & Nichols 2003), assuming that species detections were often generated by multiple individuals given the sampling design. All sets of models incorporated similar covariates for the observational and ecological processes, with spatial variation defined on the same raster landscape. We evaluate the use of occupancy as a proxy for density and discuss design-based approaches that can improve the approximation for large-scale monitoring programs.

## Materials and methods

### STUDY AREA AND SAMPLING

Our study area spanned all of western New York, USA, encompassing a 70,096 km^2^ region comprised mostly of forest and agriculture (Fuller, Linden & Royle 2016). As with other temperate forests of eastern North America, this region was historically occupied by fisher until extirpation in the early 1900s (Powell & Zielinski 1994; Lewis, Powell & Zielinski 2012). The region was delineated by 13 wildlife management unit (WMU) aggregates, 8 of which (totaling 73% of the study area) have been closed to fisher harvest since 1949 while the remaining 5 WMU’s, located near remnant and reintroduced fisher populations in the Adirondack and Catskill Mountains, have had regulated trapping seasons for >20 years (Fig. S1.1; Fuller, Linden & Royle 2016).

Our study design required a discrete representation of the landscape to define sampling units that could be surveyed for fisher and to quantify landscape attributes that might be associated with variation in fisher occurrence and density. Additional details are described in Fuller, Linden and Royle (2016). We divided the study area into a grid having 4,400 cells with a resolution of 15 km^2^, chosen to match the territory size of a female fisher (Arthur, Krohn & Gilbert 1989; Powell & Zielinski 1994). This design was intended to theoretically restrict: 1) the number of individuals within a grid cell and; 2) the number of grid cells overlapped by any given individual. The true maximum for each were unknown and would have depended on differences in movement between sexes, the amount of inter- and intrasexual overlap of territories, and the configuration of sampled grid cells. We selected a subset of available grid cells using a stratified random approach with clustering (≥ 3 neighboring cells) to accommodate field logistics. The initial sampling year in 2013 was restricted to grid cells with >60% forest cover, while sampling in 2014 and 2015 included grid cells across a broader range of forest cover values selected in proportion to landscape availability. The number of grid cells sampled in each year was 300 in 2013 and 608 in both 2014 and 2015 (Figs. S1.2–S1.4). Across all years, 826 unique grid cells were sampled, with replicated sampling across 2 and 3 years for 423 and 129 grid cells, respectively.

Sampling stations were located as close to the grid cell center as possible and consisted of a baited trap with hair snares to capture genetic samples and a remote camera for photographing species encounters. Stations were baited with beaver *Castor canadensis* meat attached to a tree and surrounded by 9 gun brushes, positioned 1–2 meters above the ground, with an infrared camera pointed at the bait tree from a location <5 meters away. Sampling occurred between January and March of each year, during which active stations were visited approximately weekly to collect hair samples (stored with silica desiccant) and replace bait as needed; 4 visits were made to each active station after initial setup in 2013, and 3 visits in 2014 and 2015.

Genetic samples were defined as a cluster of ≥5 hair follicles at a single gun brush. To reduce costs, we processed a subset of the samples for genetic data, ensuring that every site-visit combination that yielded fisher hair was included. Each processed sample had DNA extracted for species identification, molecular sexing, and microsatellite genotyping using fluorescent fragment analysis and DNA sequencers at the Cornell University Institute of Biotechnology (Ithaca, NY, USA). Genetic methods are detailed in Appendix S2.

### OCCUPANCY MODEL

We fit a single-season site occupancy model (MacKenzie *et al*. 2002) to estimate the probability of fisher occurrence in grid cells using species detections from the camera data. The model used here was described in Fuller, Linden and Royle (2016) and derived from the model selection process they used to identify important sources of variation in probabilities of detection and occupancy. Since the focus was on spatial variation in occurrence, each grid cell × year combination was considered a distinct site. Fisher detection, y_*jk*_, at site *j* during survey *k* was considered a Bernoulli random variable:

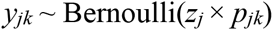

where *z_j_* is the site-specific latent occurrence state indicating whether a site is occupied (*z_j_* = 1) or not (*z_j_* = 0), and *p_jk_* is the site- and survey-specific detection probability, Pr(*y_jk_* = 1 | *z_j_* = 1). We considered each latent occurrence state a Bernoulli random variable:

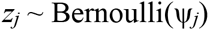

where ψ_*j*_ is the site-specific probability of occurrence, Pr(*z_j_* = 1). We used logit-link functions for each probability to examine covariates that varied by site or survey. Following the top-ranked model structure from Fuller, Linden and Royle (2016), our detection probability model was a year-specific quadratic function of ordinal date (mean of the survey week) with an effect to account for increased detection after the first survey occasion (i.e., *k* = 1 vs. *k* >1). The model for detection probability was therefore:

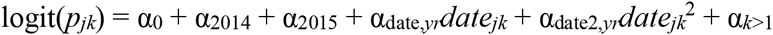

where α_2014_ and α_2015_ are additive year effects depending on when site *j* was surveyed; α_date,*yr*_ and α_date2,*yr*_ are the year-specific relationships with survey ordinal date; and α_*k*>1_ is the effect of when survey *k* >1. Our logit-linear model for occupancy included year and the 2 landscape covariates identified by model selection to be important predictors, proportion of coniferous-mixed forest and road density (Fuller, Linden & Royle 2016):

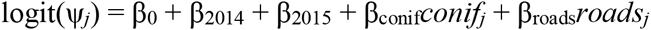

Similar to the detection probability model, β_2014_ and β_2015_ depend on the year for site *j*. The landscape covariates were calculated for each site using freely available GIS data, including the 30-m resolution National Land Cover Database (Fuller, Linden & Royle 2016). We used square-root and natural-log transformations for *conif_j_* and *roads_j_*, respectively, before scaling each to have zero mean and unit variances.

### ROYLE-NICHOLS MODEL

We fit a Royle-Nichols (RN) model (Royle & Nichols 2003) to the species detection data to examine whether heterogeneity in detection could be attributed to variation in site abundance. Our sampling design had used prior knowledge of female fisher movement to define sites, yet overlapping male and female territories or variation in individual movement could lead to sites being used by multiple individuals. Additionally, the RN model generates estimates of abundance or density using the same type of data collected for occupancy estimation, potentially providing another tool for species monitoring that does not require individual identification.

We used the same data structure described earlier for the occupancy model, with sites defined as grid cell × year combinations. Fisher detections, *y_jk_*, were modeled as Bernoulli random variables such that:

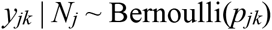

where *N_j_* is the latent site abundance and *p_jk_* is the species detection probability. Importantly, species detection probability was a function of *N_j_*:

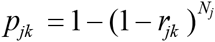

Here, *r_jk_* is the per-individual detection probability. The state process model assumed that site abundance was a Poisson-distributed random variable with mean λ_*j*_:

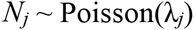

We used the same linear models to describe variation in both the observation and state process models using appropriate link functions on the relevant parameters:

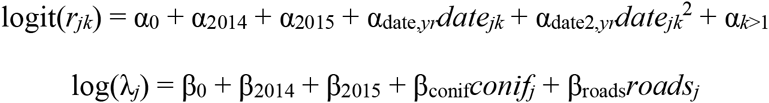

### SPATIAL CAPTURE-RECAPTURE MODEL

We used SCR (Borchers & Efford 2008; Royle & Young 2008) to model the individual encounter data generated by the genotyped hair samples and predict fisher density within the grid cells in our landscape. A standard SCR model uses the spatial distributions of individual encounters at trap locations to jointly estimate the number and location of latent activity centers (representing population size and individual distribution) and trap- and individual-specific encounter probabilities. We assumed that the encounter process for individuals exhibited similar temporal variation to that identified in the occupancy model, and that fisher density potentially varied according to the same landscape attributes influencing occurrence.

We modeled the encounter histories, y_*ijk*_, for individual *i* at trap *j* on survey *k* as Bernoulli random variables that depended on the location of the individuals latent activity center **s**_*i*_ = (s_*i*1_, s_*i*2_), such that Pr(y_*ijk*_ = 1 | **s**_*i*_) = *p_ijk_*. Importantly, encounter probability was a decreasing function of the Euclidean distance, *d_ij_*, between activity center, **s**_*i*_, and the location for trap *j*:

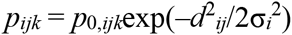

Here, *p*_0,*ijk*_ is the encounter probability when *d_ij_* = 0 while σ_*i*_ is the scale parameter of the half-normal distance function. Both parameters were made functions of covariates:

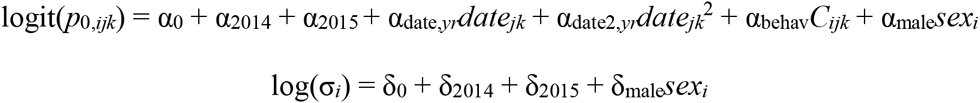

The effects for year (α_2014_, α_2014_) and ordinal date (α_date,*yr*_, α_date2,*yr*_) were similar to those for species detection probability; the survey occasion effect was replaced by a trap-specific behavioral response (α_behav_), where *C_ijk_* = 1 for all *k* after the initial encounter of individual *i* at trap *j*, and 0 otherwise. The model for σ_*i*_ also included year effects, and both models incorporated an effect for the difference between males (*sex_i_* = 1) and females (*sex_i_* = 0). We treated sex as a random variable and estimated φ_male_ = Pr(*sex_i_* = 1) using the likelihood formulation in Royle *et al*. (2015). This allowed us to estimate the sex of both un-encountered individuals and encountered individuals that could not be assigned a sex due to uncertainty in the genetic marker.

To model variation in fisher density, we used an inhomogeneous point process to describe the distribution of activity centers within our study area (Borchers & Efford 2008). We defined a discrete state space representing the possible locations of the realized point process to coincide with the 4,400 cell raster used for the occupancy model. To accommodate scales of movement (σ) that were smaller than the grid cell size, we reduced the resolution of the grid from 3.873 km × 3.873 km (15 km^2^) to 0.968 km × 0.968 km (0.938 km^2^), increasing the total number of grid cells, *G*, to 70,400. Landscape covariates were recalculated at the new resolution, though a moving-window approach was used to reflect a similar scale (15 km^2^) for the features as that assessed by the occupancy model. We modeled the expected density in a given grid cell *g* as the intensity of a point process conditional on a linear model of spatially-varying covariates, such that E(*D_g_*) = μ(*g*, **β**), where **β** are regression coefficients for the linear model (Royle *et al*. 2014). Following the model structure for occupancy, expected density was a linear function of year and the 2 landscape covariates, here on the log scale:

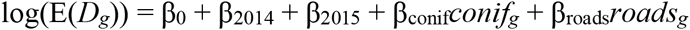

The final component of the SCR model involved defining the distribution of activity centers. Note that given the discrete state space, activity centers are now referenced by s_*i*_, a vector with the grid cell ID (*g*) for each individual, instead of the two-dimensional coordinates. For a basic SCR model having constant density, such that activity centers are distributed uniformly throughout the state space, the probability of an activity center being located in any given grid cell would be 1/*G*. Since we were modeling variation in density, the probability was a ratio of the intensity function at a given grid cell, conditional on the coefficients of the linear model and the spatial covariate values, and the summed intensity function across all grid cells:

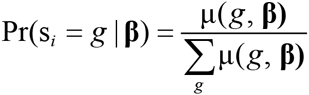

We used a Poisson-integrated likelihood approach (Borchers & Efford 2008; Royle *et al*. 2014) to evaluate the likelihood of the SCR model parameters over all possible grid cells for the activity centers.

### MODEL FITTING AND SPATIAL PREDICTIONS

All models were fit using maximum likelihood methods. For the occupancy and RN models we used the “occu” and “occuRN” functions, respectively, of the unmarked package (Fiske & Chandler 2011) in R (R Core Team 2015) to compute the likelihoods and obtain maximum likelihood estimates (MLEs). The functions used for computing the SCR likelihood and obtaining MLEs were written in R with code provided by Sutherland, Fuller and Royle (2015).

Fuller, Linden and Royle (2016) present additional information regarding model selection, relative variable importance, and goodness-of-fit in an expanded occupancy analysis of these fisher detection data from the camera trapping. For the purposes of our comparisons here, we conditioned our inferences on a single model structure for each of the model types. Following MacKenzie and Bailey (2004), we used parametric bootstrapping to assess goodness-of-fit for the occupancy and RN models and calculated an overdispersion parameter (ĉ) to compare the model types (Appendix S3). We avoided a fit assessment for the SCR model given the general lack of guidance on best practices, particularly when using maximum likelihood approaches, though we were generally less concerned with model fit given the flexibility and robustness of SCR to deviations from model assumptions (Royle *et al*. 2014). We also considered the SCR model to represent a better approximation to the actual state variable of interest (i.e., individual fisher distribution) than either model using detection-nondetection data.

We generated spatial predictions from each model for the 4,400 grid cells in the landscape of interest. For the occupancy and RN models, landscape covariates for all grid cells were transformed and then scaled using the values calculated across the surveyed grid cells. For the RN model, we generated spatial predictions of expected fisher density (#/km^2^) using E(λ_*j*_)/15, the expected abundance divided by area for a grid cell. For the SCR model, expected density (#/grid cell) was predicted across the high-resolution state space (*G* = 70,400) using the MLEs for **β**, then an aggregate mean density (#/km^2^) was calculated for each of the 4,400 cells from the original grid. Following Fuller, Linden and Royle (2016), we calculated average values for the grid cells within each of the 13 WMUs to compare how management decisions may differ between the models. Finally, we used least-squares regression to examine relationships between predictions from the models using detection-nondetection data (occupancy and RN) to predictions from the SCR model. Values were transformed to the appropriate scale (logit or log) before fitting the regressions. Hereafter, we refer to each regression according to the relationship that was modeled, where “SCR=occupancy” was log(SCR density) ~ logit(occupancy) and “SCR=RN” was log(SCR density) ~ log(RN density). We report the slope coefficient for the SCR=RN regression given that a 1:1 relationship was possible.

## Results

Cameras detected fisher at 198/289 (67%) operational traps in 2013 (11 cameras malfunctioned), 310/608 (51%) traps in 2014, and 236/608 (39%) traps in 2015 (Figures S1.2–S1.4). Hair deposits confirmed to be fisher were collected at a fraction (range: 0.32–0.45) of sites with confirmed camera captures (Table 1). Identity assignment of 178, 281, and 138 successfully genotyped hair samples resulted in 89, 165, and 90 unique individuals encountered in 2013, 2014, and 2015, respectively (Table 1); the number of spatial recaptures (individuals encountered in >1 trap) was 8, 9, and 3 in each year. Observed sex ratio across all years was approximately even (143 F, 145 M, 56 NA).

**Table 1.** Results from baited hair snares during the winters (Jan–March) of 2013–2015 used to detect fisher in western, New York.

| Year | Traps |  | Fisher specimens |  |  |
| --- | --- | --- | --- | --- | --- |
|  | Total | Fisher hair<sup>a</sup> | Total | Genotyped<sup>b</sup> | Individuals |
| 2013 | 300 | 82 | 377 | 178 | 89 |
| 2014 | 608 | 141 | 425 | 281 | 165 |
| 2015 | 608 | 76 | 455 | 138 | 90 |

<sup>a</sup> Number of unique traps with $\geq 1$ hair sample confirmed to be fisher.
<sup>b</sup> Successfully genotyped specimens defined as having $\leq 3$ missing loci of the 9 total loci used to determine individual genotypes.

Model estimates for the observation processes of the 3 model types indicated similar patterns within and between years (Table 2), including a decrease in average detection and encounter probabilities from 2013 to 2015. The regression coefficients for observation covariates were nearly identical for the occupancy and RN models, which was expected given that they used the same detection-nondetection data. The directions of most effects in the SCR model matched those for the other models; the only coefficient that differed (α_date2,2014_) was estimated near zero for each model. Across years, mean detection probability during survey 1 for the species ranged 0.16–0.49, while that for individuals ranged 0.10–0.29; for surveys >1, the ranges of mean detection probabilities increased to 0.25–0.62 and 0.16–0.41 for species and individuals, respectively. The SCR model indicated a strong local behavioral response (α_behav_ = 3.842 [SE: 0.366]), suggesting individuals were much more likely to return to a trap after an initial visit. The log-linear coefficients for σ*i* indicated that the movement scale was larger in the later years (δ_2014_ = 0.456 [SE: 0.313]; δ_2015_ = 0.811 [SE: 0.358]) and for males (δ_male_ = 0.298 [SE: 0.341]). This resulted in mean σ_*i*_ estimates that ranged 3.55–7.98 km for females and 4.78–10.76 km for males, across the years. The goodness-of-fit statistics indicated some overdispersion for the models using detection-nondetection data (Appendix S3), more so for the occupancy model (ĉ = 2.78) than for the RN model (ĉ = 1.59).

**Table 2.** Parameter estimates from the observation process components of the occupancy, Royle-Nichols (RN), and spatial capture-recapture (SCR) models. The parameters describe the logit-linear model (**α**) of trap-specific encounter probability for individuals (RN, SCR) or species (occupancy), and the log-linear model (**δ**) of the half-normal distance function in SCR.

| Parameter | Occupancy |  | Royle-Nichols |  | Spatial capture-recapture |  |
| --- | --- | --- | --- | --- | --- | --- |
|  | Mean | SE | Mean | SE | Mean | SE |
| $\alpha_0$ | -0.056 | 0.131 | -0.886 | 0.168 | -4.475 | 0.457 |
| $\alpha_{2014}$ | -0.538 | 0.165 | -0.498 | 0.226 | -0.383 | 0.416 |
| $\alpha_{2015}$ | -1.579 | 0.192 | -1.284 | 0.258 | -1.196 | 0.518 |
| $\alpha_{\text{date},2013}$ | 0.147 | 0.030 | 0.139 | 0.030 | 0.644 | 0.150 |
| $\alpha_{\text{date},2014}$ | 0.113 | 0.023 | 0.112 | 0.022 | 0.166 | 0.081 |
| $\alpha_{\text{date},2015}$ | -0.020 | 0.022 | -0.021 | 0.022 | -0.107 | 0.087 |
| $\alpha_{\text{date}2,2013}$ | -0.031 | 0.008 | -0.028 | 0.007 | -0.317 | 0.101 |
| $\alpha_{\text{date}2,2014}$ | 0.002 | 0.007 | 0.004 | 0.007 | -0.027 | 0.068 |
| $\alpha_{\text{date}2,2015}$ | 0.049 | 0.007 | 0.044 | 0.007 | 0.264 | 0.083 |
| $\alpha_{k>1}$ | 0.548 | 0.088 | 0.512 | 0.081 | — | — |
| $\alpha_{\text{behav}}$ | — | — | — | — | 3.842 | 0.366 |
| $\alpha_{\text{male}}$ | — | — | — | — | -0.425 | 0.420 |
| $\delta_0$ | — | — | — | — | 1.267 | 0.242 |
| $\delta_{2014}$ | — | — | — | — | 0.456 | 0.313 |
| $\delta_{2015}$ | — | — | — | — | 0.811 | 0.358 |
| $\delta_{\text{male}}$ | — | — | — | — | 0.298 | 0.341 |

The relationships between the landscape covariates and the ecological processes for each model type were largely consistent (Table 3), with effects of coniferous-mixed forest proportion being significantly positive and those of road density being significantly negative. There was little support for differences in occupancy and abundance across the years. Mean fisher occupancy per 15-km^2^ grid cell was relatively high (0.69 [95% CI: 0.61–0.76]), while mean fisher density (#/km^2^) was relatively low for both the RN model (0.09 [95% CI: 0.07–0.11]) and the SCR model (0.05 [95% CI: 0.02–0.10]). The probability of being male was estimated as 0.47 [95% CI: 0.23–0.73] in the SCR model, suggesting a nearly even sex ratio.

**Table 3.** Parameter estimates from the ecological process components of the occupancy, Royle-Nichols (RN), and spatial capture-recapture (SCR) models. The parameters describe the linear model (**β**) of species occurrence probability on the logit scale (occupancy) or abundance on the log scale (RN, SCR). The intercept represents the mean for a 15-km^2^ grid cell (occupancy, RN) or a 0.938-km^2^ grid cell (SCR). The SCR model includes the logit-scale probability of being male (φ_male_).

| Parameter | Occupancy |  | Royle-Nichols |  | Spatial capture-recapture |  |
| --- | --- | --- | --- | --- | --- | --- |
|  | Mean | SE | Mean | SE | Mean | SE |
| $\beta_0$ | 0.789 | 0.172 | 0.281 | 0.109 | -3.159 | 0.407 |
| $\beta_{2014}$ | -0.382 | 0.209 | -0.143 | 0.147 | -0.250 | 0.511 |
| $\beta_{2015}$ | -0.172 | 0.268 | -0.197 | 0.176 | -1.004 | 0.562 |
| $\beta_{\text{conif}}$ | 0.435 | 0.081 | 0.223 | 0.037 | 0.583 | 0.088 |
| $\beta_{\text{roads}}$ | -0.302 | 0.094 | -0.103 | 0.043 | -1.174 | 0.282 |
| $\phi_{\text{male}}$ | — | — | — | — | -0.120 | 0.556 |

The spatial predictions indicated similar patterns of variation (Figure 1), which was expected given the consistency in estimated relationships with the landscape covariates. The SCR=occupancy and SCR=RN regressions both had R^2^ values >0.94 (Figure S1.5), though the slope coefficient for the SCR=RN regression was 2.670 [SE: 0.010]. The slope >1 was consistent with the smaller estimates for β_conif_ and β_roads_ in the RN model than in the SCR model (Table 3) and indicated the strengths of association between fisher density and the landscape covariates were reduced in the RN model. The spatial distribution of residuals further illustrated how the SCR=occupancy and SCR=RN regressions generally overpredicted in areas of low density, and underpredicted in areas of high density (Figure 1B–C). Mean values across WMUs exhibited a strong correlation between predictions from the SCR model and those from the occupancy and RN models (Figure 2), indicating largely similar inferences regarding variation in fisher distribution and density at the management-unit level.

**Figure 1.**
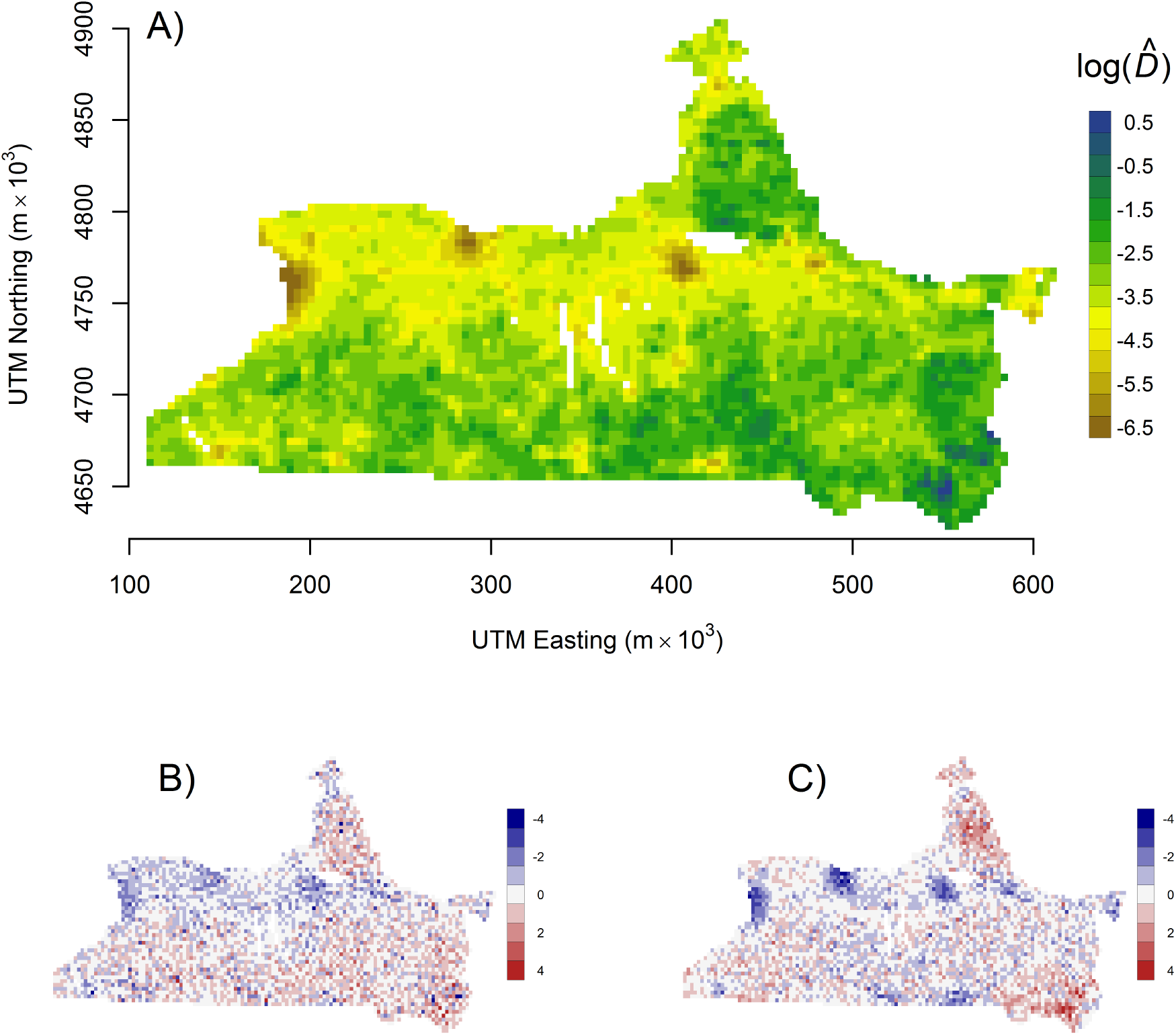
Spatial predictions from the models of fisher occupancy and abundance in western New York, including: A) expected fisher density (#/km^2^) on the log scale as predicted by the SCR model; B) standardized residuals from the SCR=occupancy regression; and C) standardized residuals from the SCR=RN regression. Blue values in (B,C) represent grid cells where the detection-nondetection data overpredicted density, while red values represent underpredictions.

**Figure 2.**
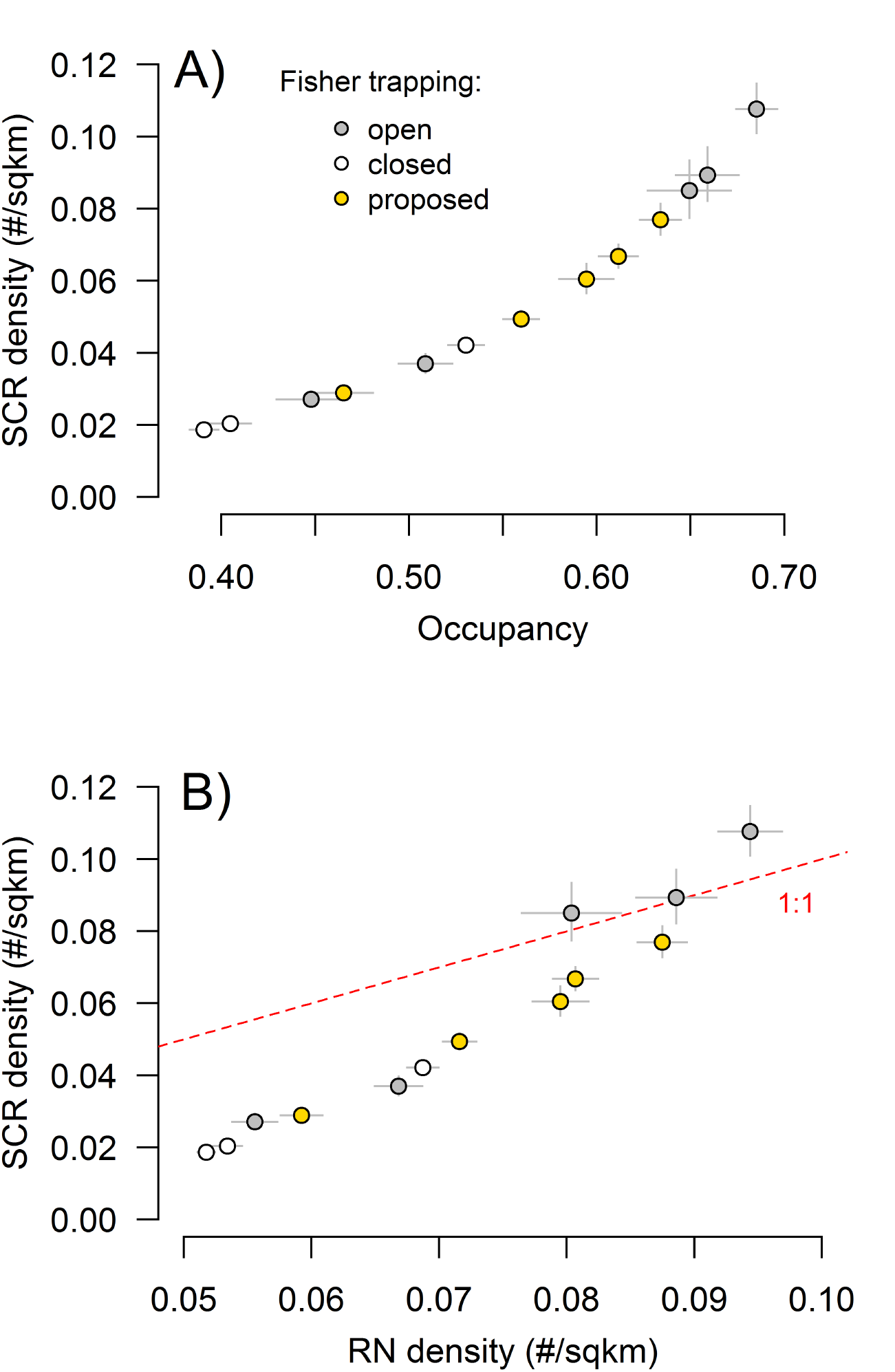
Mean (± 2 SE) predicted occupancy and density for wildlife management unit (WMU) aggregates in western New York, 2013–2015, comparing estimates from the SCR model to those from the occupancy model (A) and the RN model (B). The trapping status for each WMU indicates whether fisher harvest is open (gray), closed (white), or being proposed for opening (yellow). Red line in (B) indicates 1:1 relationship.

## Discussion

An understanding of the statistical and ecological relationships between species occupancy and density can improve the design of monitoring programs that aim to make inferences on wildlife populations at large scales. Complete knowledge on the distribution of individuals would provide the necessary information for management and conservation, but such data almost never exist, particularly for wide-ranging terrestrial species such as carnivores. Therefore, the distribution of individuals must be sampled and statistically summarized, with limitations for each step being determined by the selected study design and the ecology of the focal species. Sampling a collection of sites for species occurrence is typically easier and less cost prohibitive at large scales than sampling and identifying individuals, yet the resulting data summary may not provide an adequate approximation to the information of interest. We presented empirical evidence that models of occupancy and density can generate similar predictions and management recommendations when species movement ecology is considered in the sampling design, even when some modeling assumptions are violated.

Our study currently represents one of the largest applications of spatial capture-recapture modeling in terms of both landscape extent and coverage. Obbard, Howe and Kyle (2010) have the only comparable extent (across Ontario, Canada) for an SCR study, but their sampling was of distinct populations and did not involve contiguous landscape predictions. A benefit to our comprehensive effort was that spatially-explicit density estimates could be used to evaluate the ability of occupancy models to guide wildlife management decisions at regional scales (Clare, Anderson & Macfarland 2015; Fuller, Linden & Royle 2016). Our evaluation suggests that detection-nondetection data can be a useful tool for indexing density and addressing large-scale monitoring needs under certain conditions. The estimates of average fisher occupancy and density were highly correlated at the WMU scale (Figure 2) and would likely lead to similar management decisions regarding expanded harvest opportunities (Fuller, Linden & Royle 2016).

The correspondence among model types was largely due to the strong associations between the estimated state variables and the two landscape attributes that exhibited wide variation across the region. The positive effect of coniferous-mixed forest and negative effect of road density are consistent with previous knowledge on fisher ecology (Powell 1993; Powell & Zielinski 1994). Interestingly, the models using detection-nondetection data had relatively smaller effect sizes for each covariate, though this result could have been expected. The occupancy model involves a logistic regression of latent occurrence, a binary random variable, yet the number of individual fisher occurring within or using a grid cell could be >1. Therefore, the coefficients in the logit-linear model should underestimate the effects of covariates if variation exists among grid cells with ≥1 individual. Estimates from the SCR model indicated that individual movement for both sexes was large enough to encompass multiple grid cells and, combined with intersexual territory overlap (Powell 1993), would have resulted in many sampled grid cells having >1 individual in areas of relatively high density. The RN model was specifically designed to deal with abundance-induced heterogeneity in detection probability (Royle & Nichols 2003) and had a lower estimate of overdispersion than the occupancy model, suggesting a potentially superior fit. Both models of detection-nondetection data struggled to explain the number of observed detection histories that were all 1s or all 0s (Appendix S3), suggesting unmodeled heterogeneity. Given sex-specific differences in movement and individual differences in the location of activity centers, the assumption of equal per-individual detection probability for the RN model was clearly violated, as was the assumption of independence between sites for the occupancy model. Despite these limitations, each model of detection-nondetection data served as an adequate index for density, as estimated by SCR.

Additional model complexity may have improved our use of the detection-nondetection data, for example, by incorporating a spatial dependence structure (Johnson *et al*. 2013). The size of our grid cells in comparison to the observed animal movement and the clustered pattern of sampled cells may have warranted some type of autoregressive function. Johnson *et al*. (2013) present an approach that is computationally efficient for large landscapes and, importantly, addresses concerns with possible confounding between latent spatial effects and landscape or habitat covariates (Hodges & Reich 2010). For many species, this approach may be particularly useful when ecological or observational processes exhibit spatial correlation and have the potential to affect inferences. In our application of occupancy modeling, the latent spatial process that likely caused overdispersion problems was related to individual distribution – the very state variable that is estimated by SCR. Depending on the scale and extent of the study, SCR represents a more comprehensive approach for making inferences about species distribution in continuous landscapes than that which can be estimated by detection-nondetection data (Efford & Dawson 2012).

An ideal monitoring program for wide-ranging, low-density species such as carnivores involves a study design that can collect individual encounters across a large landscape and allow for fitting the data to spatial capture-recapture models. Unfortunately, the associated costs and logistical difficulties will limit the application of such designs in many situations (Efford & Dawson 2012; Ellis, Ivan & Schwartz 2014), particularly when the species cannot be easily identified using distinguishable individual features (Sollmann *et al*. 2011; Clare, Anderson & Macfarland 2015). While noninvasive genetic sampling can solve the identity problem for species without unique markings, a drawback is the often low amplification rates due to poor quality DNA (e.g., from hair follicles), resulting in data with fewer useable individual encounters than the species detections that could be obtained by other means. Integrated approaches allow opportunities to calibrate inferences from detection-nondetection data by periodically including more expensive or intensive sampling to obtain individual encounters (Chandler & Clark 2014), and may represent the best compromise for designing robust monitoring programs that can make inferences across time and space. When occupancy alone is chosen as a cost-effective state variable for monitoring, simulation and sensitivity analyses should be used to understand how inferences from detection-nondetection data will be affected by aspects of study design and species ecology (Ellis *et al*. 2015).

## Acknowledgments

We thank the following NYSDEC staff for coordinating and conducting field surveys: K. Baginski, M. Clark, E. Duffy, L. Durfey, R. Holevinski, A. MacDuff, M. Putnam, A. Rothrock, B. Schara, and S. Smith. We thank B. Swift, M. Schiavone, and P. Jensen for project support and H. Borchardt-Wier for genotyping. We thank R. Holevinski for assistance in obtaining relevant GIS databases and for helping to coordinate field efforts. This work was supported in part by Federal Aid in Wildlife Restoration Grant W-173-G. Any use of trade, firm, or product names is for descriptive purposes only and does not imply endorsement by the U.S. Government.

## Data accessibility

Data will be archived with Dryad Digital Repository.

## Appendix S1. Additional figures

**Figure S1.1.**
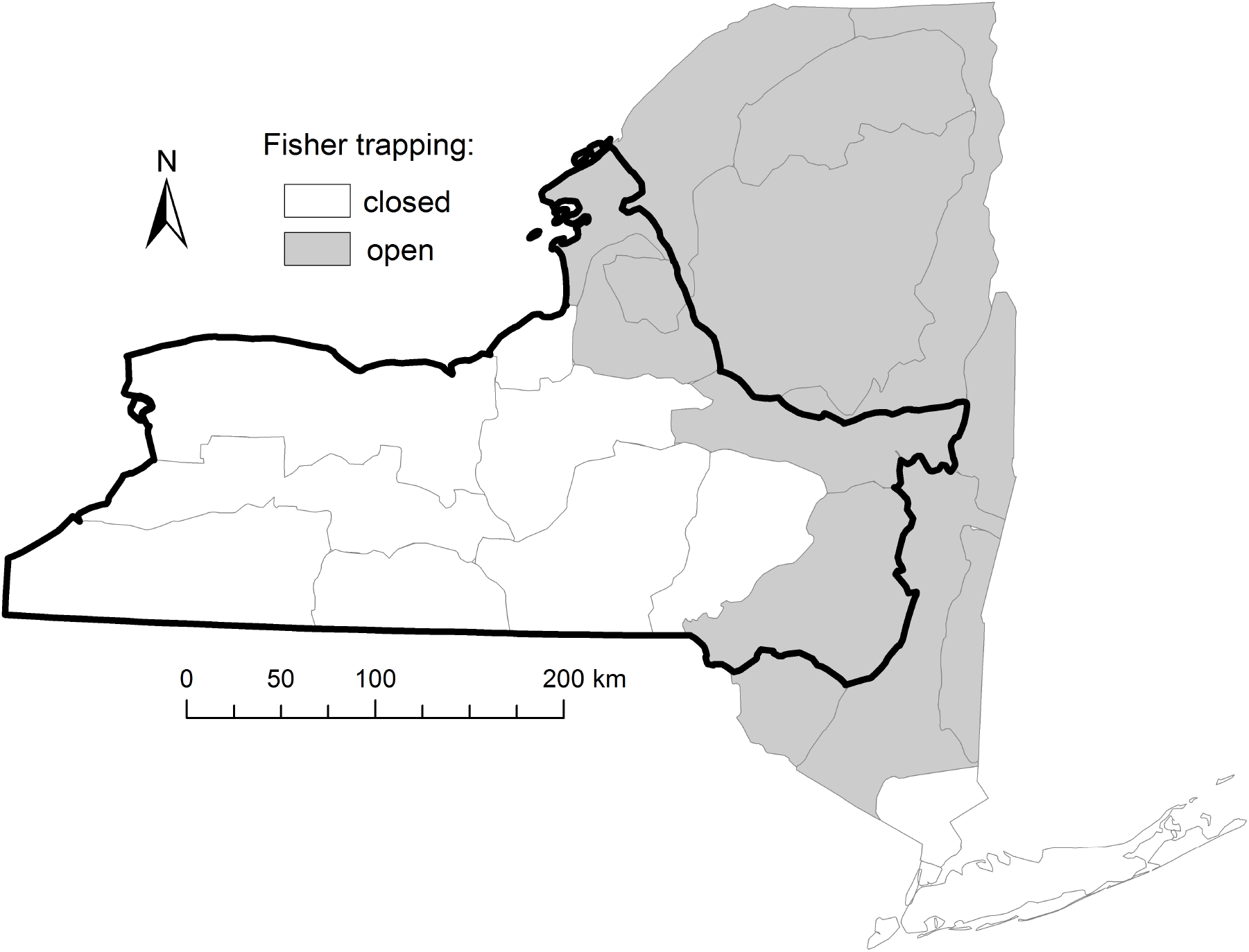
Map of study area in western New York, USA, outlined in bold with delineations for aggregated wildlife management units that were open (gray) and closed (white) to fisher trapping as recently as 2016.

**Figure S1.2.**
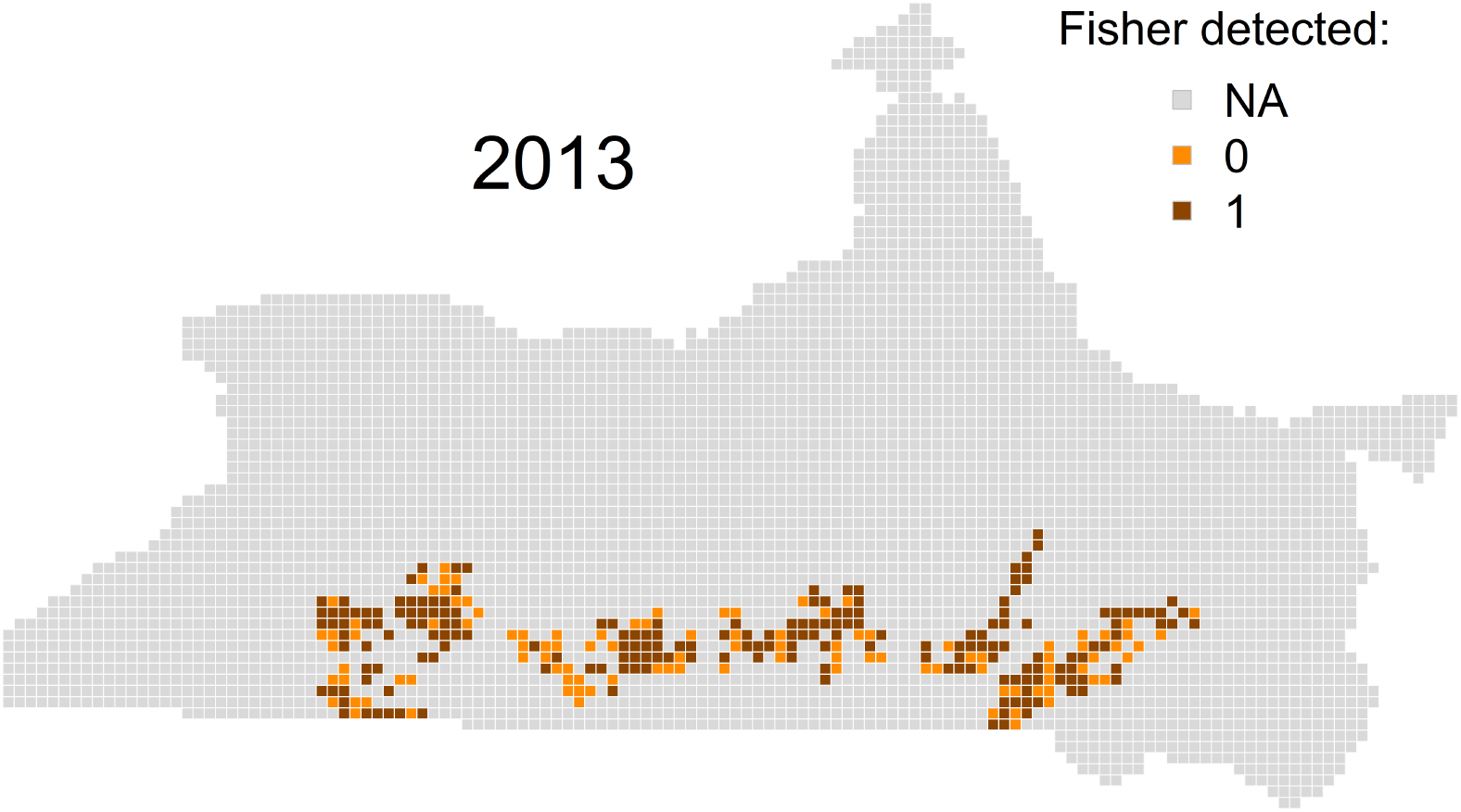
Grid cells noninvasively sampled for fisher in 2013 in western New York, USA, that resulted in 0 detections (light orange) or ≥1 detections (dark orange).

**Figure S1.3.**
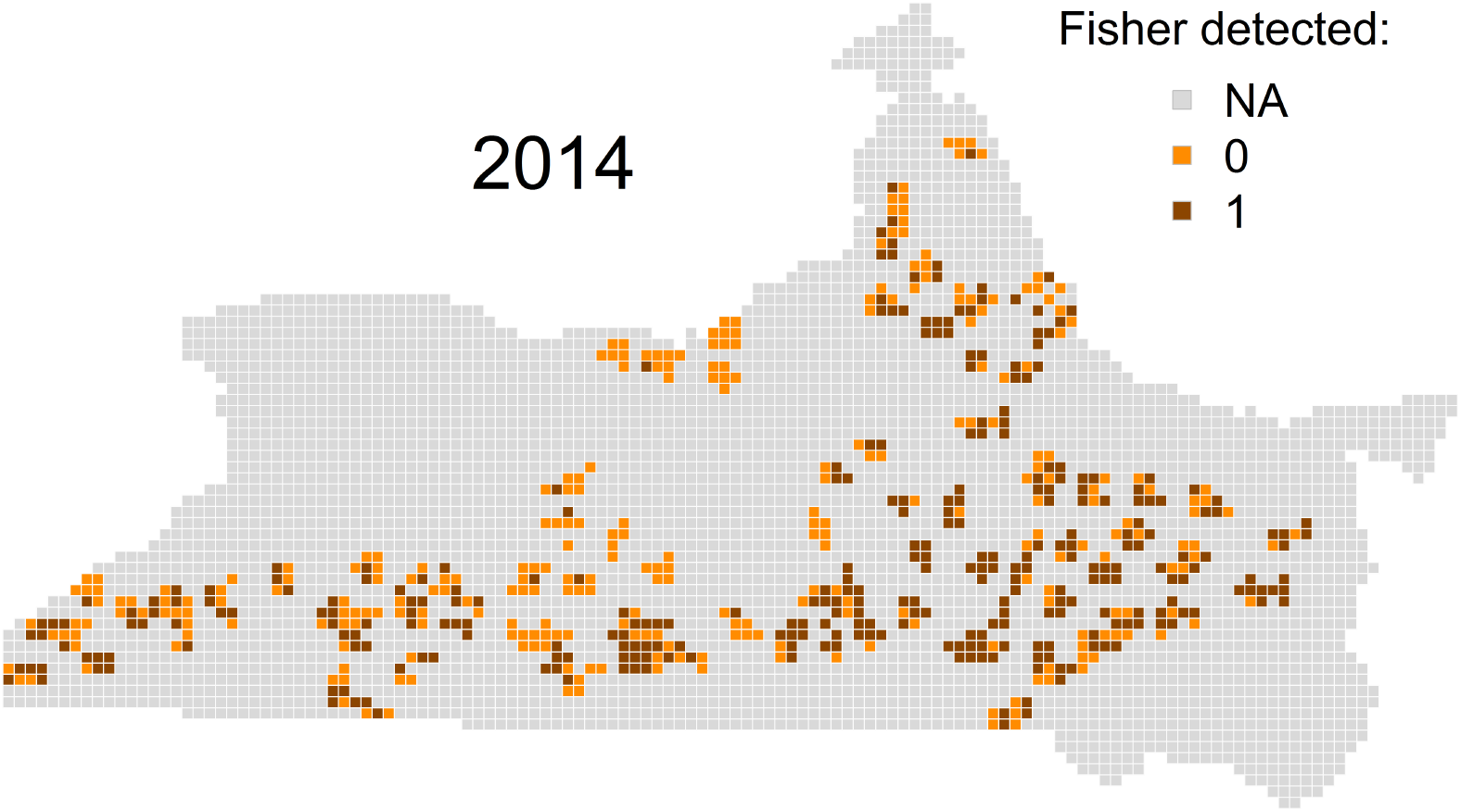
Grid cells noninvasively sampled for fisher in 2014 in western New York, USA, that resulted in 0 detections (light orange) or ≥1 detections (dark orange).

**Figure S1.4.**
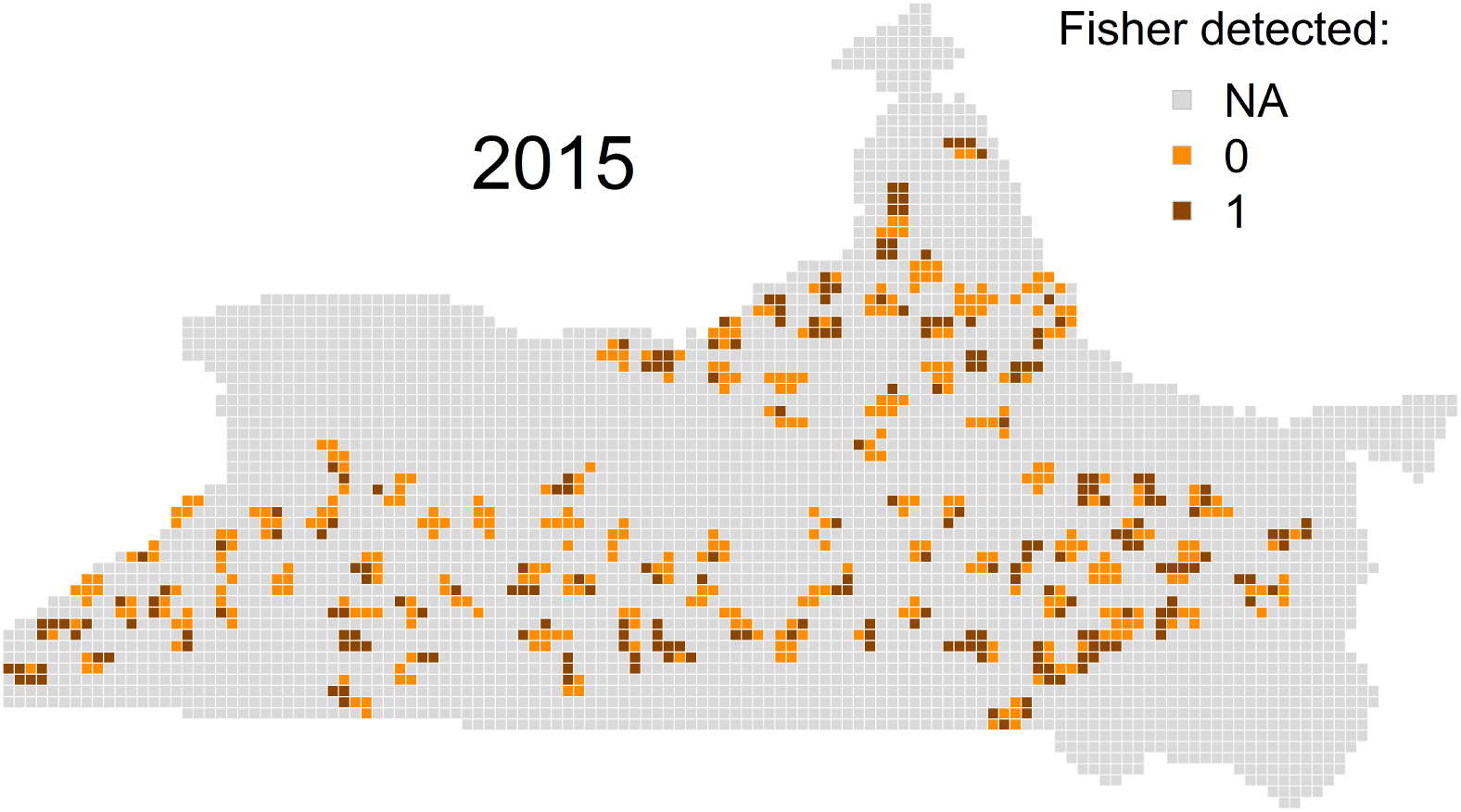
Grid cells noninvasively sampled for fisher in 2015 in western New York, USA, that resulted in 0 detections (light orange) or ≥1 detections (dark orange).

**Figure S1.5.**
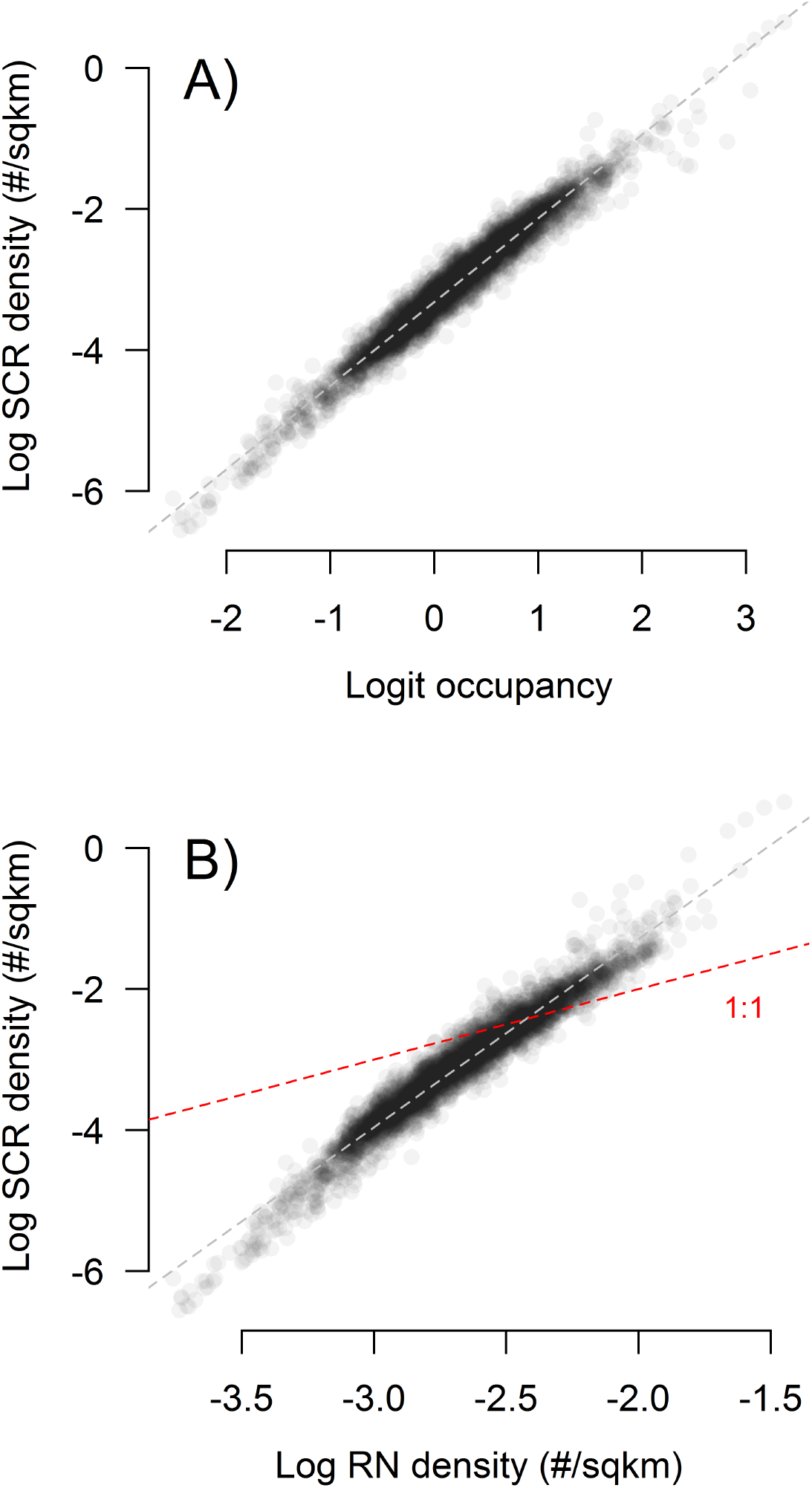
Grid-cell relationships between link-scale predicted values from the models for detection-nondetection data, including the occupancy (A) and Royle-Nichols (B) models, and predicted values from the spatial capture recapture model. The red dashed line in (B) represents a 1:1 relationship between the log-scale density predictions.

## Appendix S2. Technical details of fisher genetic methods

Genotyping included mitochondrial DNA for species determination, a Y chromosome marker for sex determination, and autosomal microsatellite markers for individual identification. Genotyping for 2013 and 2014 samples initially targeted the specimen from each unique visit to a trap site (hereafter, site-visit) that had the greatest number of hairs with follicles. It was required that samples had ≥5 hairs. Subsequently, we increased sampling by processing samples from an additional gun brush at site-visits where the first did not amplify DNA (2013) or for every site-visit with ≥5 hairs on a second gun brush (2014). Total number of samples collected in 2015 were fewer than in previous years, so no subsampling was conducted. DNA extraction used DNeasy 96 plates (Qiagen) following the manufacturer’s protocol for tissue, except for addition of 20 ul of 1M DTT at the ATL/Proteinase K step.

Samples were determined to be fisher based on NCBI blastn comparisons of a segment of mitochondrial D-loop amplified and sequenced with primers designed to work for mustelid species. We designed general mustelid primers directed at mitochondrial tRNA-proline and control region, 15409Md-F: 5’-CCCAAAGCTGAYATTCTAA-3’ and 15657Md-R: 5’-TTGMTGGTTTCTCGAGGC-3’ (names based on Genbank Accession HM106322.1, *Neovison vison* complete mitochondrion genome). The PCR reactions were 20 ul total volume and contained 1 ul DNA, 0.2 mg/ml BSA, 0.1 mM each dNTP, 2.5 mM MgCl2, 0.2 uM each primer, 0.4 U taq DNA polymerase and 1x vendor’s PCR buffer (Invitrogen). PCR cycling started with 95° C for 2 min, then 31 cycles of 95° C for 1 min, 57° C for 1 min, 72° C for 1 min, followed by 72° C for 5 min. Amplicons were cleaned with ExoSAP-IT (Fisher Scientific) and sequenced with BigDye 3.1 (Applied Biosystems) on one strand. Resulting sequences were blasted to the NCBI GenBank database and even partial sequence provided unambiguous identification of species.

Sex was assigned based on co-amplification of an intron segment in the Y-linked DBY7 gene and a mitochondrial DNA internal positive control. Amplification of the Y chromosome was from intron 7 of the DBY7 gene using primers DBY7.2F: 5’-TTAGTTGGGACCTTTCTTTCTAAACAG-3’ and DBY7.2R: 5’-TGGTATCGGGTCCCACAT-3’. PCR reactions included 2 ul of DNA template in a 20 ul total volume with 0.2 mg/ml BSA, 0.2 mM dNTPs, 2.5 mM MgCl2, 0.5 uM each DBY7.2 primer, 1 unit of Platinum hot start taq with 1x vendor’s buffer (Invitrogen). In addition, 0.05 ul each of mitochondrial primers 15409MdF and 15657MdR (see above) were used as an internal positive control (IPC). Thermocycling conditions were 95° C for 15 minutes, touchdown PCR of 95° C for 30 s, 65–55° C for 30 s, 72° C for 30 s for 20 cycles, dropping 0.5° C per cycle, followed by 20 cycles of 95° C for 30 s, 55° C for 30 s, 72° C for 30 s, with a final extension of 72° C for 5 min. Amplicons from at least 3 replicate PCRs were scored in 2.5% agarose gels with ethidium bromide staining. Specimens were scored as female if only the IPC amplified (fragment size = 250 bp) for all 3 replicates, male if the DBY7.2 fragment (size =190) amplified in at least 2 replicates, or unknown if the gel patterns were inconsistent across replicates.

Nine previously characterized microsatellite loci were amplified in three multiplexes: GGU101, MP0055, MP0084, MP0100, MP0182 (Jones *et al*. 2007), MA1 (Davis & Strobeck 1998), MVIS072 (Fleming *et al*. 1999), LUT604 (Dallas & Piertney 1998), and RIO20 (Beheler *et al*. 2005). Forward primers were fluorescently labeled and reverse primers were pigtailed with the sequence GTTTCTT to reduce incomplete adenylation in PCR (Table S2.1). PCR was initially conducted for 3–4 replicates on each specimen. Based on a consensus genotype from the initial replicates, specimens amplifying at <5 loci were culled from the analysis. Missing data in the remaining specimens, if any, triggered up to 3 additional replicates. Optimized PCR reactions included 2 ul DNA, 0.2 mg/ml BSA, 1.5 mM MgCl2, 0.2 mM each dNTP, 0.16 uM each primer, 0.4 U Platinum hot start taq (Invitrogen) and 1.5x vendor’s PCR buffer in 10 ul total volume. Negative and positive control samples were included in every PCR setup, and all PCR reactions were assembled in a separate pre-PCR lab, under a hood after UV treatment of materials. Thermocycling parameters for three multiplexes were 94° C for 2 min; 94° C for 30 s; 58° C for 45 s; 72° C for 45 s for 40 cycles; followed by a 30-minute final extension at 72° C.

Many samples in 2015 showed very poor amplification success rates based on Qiagen extractions, so for those that had been identified as fisher and had remaining hair (n = 108) we performed additional replicate PCRs using Phire Tissue Direct PCR Master Mix (Thermo Fisher, cat no F-170L), following the dilution protocol with 1–2 hairs. A total of 23 samples were analyzed with 3–4 replicates of both Qiagen and Phire PCR and only 1.3% allelic mismatches were found between consensus genotypes, indicating comparability of data from the two methods.

Automated calling of genotypes was done with Genemapper 4.0 (Applied Biosystems) followed by manual checking of call accuracy. A consensus genotype for each specimen was determined using the comparative multiple-tubes approach of Frantz *et al*. (2003) in which heterozygotes require two observations and homozygotes require three to be called. Ambiguous genotypes (e.g., insufficient replication accomplished) were scored as missing, and ≤3 loci were missing out of 9 in the final analyzed samples. Any replicated observation of ≥3 alleles at a locus caused the specimen to be dropped from analysis (possible contamination).

A PI(sib) threshold of 0.005 was applied to filter specimens with lower information content in their multilocus genotypes, and then this same threshold was applied pairwise (match probability) to create provisional transitive recapture clusters. The PI(sib) thresholding follows Creel *et al*. (2003) and Sethi *et al*. (2014) to remove multilocus genotypes with low confidence in distinguishing siblings. Allele frequencies for PID calculations were estimated from 2013–2015 specimens that differed from all other specimens by ≥4 allelic differences (n = 302). To combine two or more specimens into provisional recapture groups, we required that they have an observed pairwise PI(sib) (match probability based on all identical loci) <0.005. Subsequent error-informed manual adjustments to the provisional PID recapture groups were based on the higher information content in recapture groups. A recapture group was combined with a provisional singleton specimen if the number of locus mismatches to the group consensus was ≤2. Provisional groups based solely on the PI(sib) match probability threshold were split (one specimen removed) if they contained specimens with false allele differences to the consensus in ≥2 loci and allelic dropout in ≥1 additional locus, or any combination of ≥4 locus differences.

**Table S2.1:**
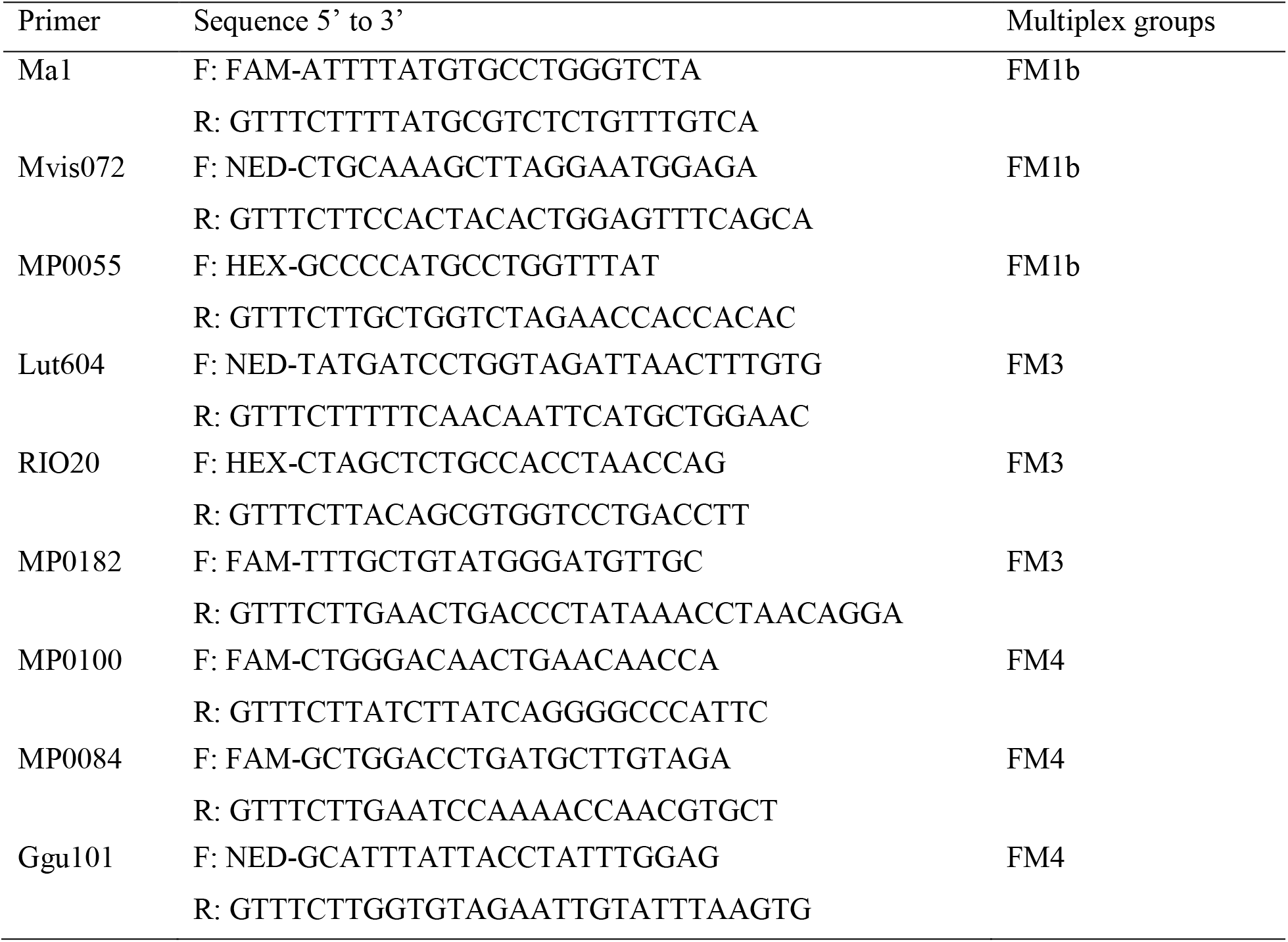
Primer sequences and multiplex groups for 9 microsatellite loci used in the NY fisher study.

## Appendix S3. Goodness-of-fit for models of detection-nondetection data

We assessed the goodness-of-fit for the two models using detection-nondetection data (occupancy and Royle-Nichols) according to methods described by MacKenzie and Bailey (2004) for site occupancy models. The approach involves calculating a Pearson’s chi-square fit statistic to the observed and expected frequencies of detection histories for a given model. Parametric bootstrapping is used to approximate the distribution of the fit statistic by simulation, accounting for likely deviations from the theoretical distribution under small sample sizes with low expected frequencies. An overdispersion parameter (ĉ) is calculated as the ratio of the observed fit statistic to the mean of the simulated distribution, with values >1 indicating overdispersion (variance > mean).

For the occupancy model, we used the “mb.gof.test” function in the AICcmodavg package (Mazerolle 2015) in R (R Core Team 2015). This function can handle occupancy models produced by the “occu” function in unmarked (Fiske & Chandler 2011) to calculate the observed and expected frequencies of the detection histories. For the RN model that we fit using the “occuRN” function we needed to modify the source code of the fit test to accommodate the altered likelihood structure. This additional functionality may be added to “mb.gof.test” in future updates (M. Marzerolle, personal communication). We simulated 1,000 bootstrap samples for each fit assessment.

The goodness-of-fit comparison indicated that the observed detection history data were overdispersed for both models, more so for the occupancy model (ĉ = 2.78) than the RN model (ĉ = 1.61). The fit of individual detection histories differed markedly between the models (Table S3.1), likely due to the additional uncertainty introduced by having to integrate over possible values of *N* given an observed detection history for the RN model. Despite individual detection histories having larger discrepancies between observed and expected frequencies for the RN model, the observed data appeared to be a closer realization to the expected distribution under the RN model, thus indicating a lower ĉ estimate than that for occupancy. More simulation testing is necessary to fully explore the adequacy of such a fit assessment for the RN model (MacKenzie & Bailey 2004).

**Table S3.1:** Distribution of observed and expected frequencies under both the occupancy and Royle-Nichols models for the most common cohorts of detection histories (>90%) from the detection-nondetection data collected at camera traps in the NY fisher study.

| History | Cohort | Observed | Occupancy |  | Royle-Nichols |  |
| --- | --- | --- | --- | --- | --- | --- |
|  |  |  | Expected | Chi-square | Expected | Chi-square |
| 0000 | 0 | 74 | 69.53 | 0.29 | 33.57 | 48.69 |
| 0001 | 0 | 18 | 12.09 | 2.89 | 23.35 | 1.23 |
| 0010 | 0 | 11 | 10.86 | 0.00 | 22.09 | 5.57 |
| 0011 | 0 | 7 | 14.77 | 4.09 | 17.04 | 5.92 |
| 0100 | 0 | 15 | 9.62 | 3.01 | 20.43 | 1.44 |
| 0101 | 0 | 5 | 13.20 | 5.10 | 15.80 | 7.38 |
| 0110 | 0 | 5 | 13.04 | 4.96 | 15.73 | 7.32 |
| 0111 | 0 | 17 | 19.00 | 0.21 | 13.53 | 0.89 |
| 1000 | 0 | 13 | 4.85 | 13.72 | 11.91 | 0.10 |
| 1001 | 0 | 7 | 6.66 | 0.02 | 9.17 | 0.51 |
| 1010 | 0 | 6 | 6.84 | 0.10 | 9.37 | 1.21 |
| 1011 | 0 | 5 | 9.89 | 2.42 | 7.96 | 1.10 |
| 1100 | 0 | 8 | 6.84 | 0.20 | 9.28 | 0.18 |
| 1101 | 0 | 5 | 9.75 | 2.31 | 7.84 | 1.03 |
| 1110 | 0 | 3 | 10.51 | 5.37 | 8.24 | 3.33 |
| 1111 | 0 | 34 | 15.54 | 21.92 | 7.70 | 89.91 |
| 000NA | 1 | 627 | 619.33 | 0.09 | 517.92 | 22.97 |
| 001NA | 1 | 116 | 113.25 | 0.07 | 175.62 | 20.24 |
| 010NA | 1 | 105 | 103.93 | 0.01 | 164.37 | 21.45 |
| 011NA | 1 | 96 | 94.02 | 0.04 | 71.95 | 8.04 |
| 100NA | 1 | 61 | 61.81 | 0.01 | 105.82 | 18.98 |
| 101NA | 1 | 21 | 51.90 | 18.40 | 44.48 | 12.39 |
| 110NA | 1 | 37 | 50.66 | 3.68 | 43.31 | 0.92 |
| 111NA | 1 | 84 | 52.10 | 19.53 | 23.53 | 155.42 |

